# Detecting Protein-DNA Binding in Single Molecules using Antibody Guided Methylation

**DOI:** 10.1101/2023.11.20.567792

**Authors:** Apoorva Thatavarty, Naor Sagy, Michael R Erdos, Isac Lee, Jared T Simpson, Winston Timp, Francis S Collins, Daniel Z Bar

## Abstract

Characterization of DNA binding sites for specific proteins is of fundamental importance in molecular biology. It is commonly addressed experimentally by chromatin immunoprecipitation and sequencing (ChIP-seq) of bulk samples (10^3^-10^7^ cells). We have developed an alternative method that uses a Chromatin Antibody-mediated Methylating Protein (ChAMP) composed of a GpC methyltransferase fused to protein G. By tethering ChAMP to a primary antibody directed against the DNA-binding protein of interest, and selectively switching on its enzymatic activity *in situ*, we generated distinct and identifiable methylation patterns adjacent to the protein binding sites. This method is compatible with methods of single-cell methylation-detection and single molecule methylation identification. Indeed, as every binding event generates multiple nearby methylations, we were able to confidently detect protein binding in long single molecules.

Graphical abstract
(i) ChAMP is added to fix and permeabilized cells, where it binds (ii) any antibody, and upon the addition of SAM, methylates nearby GpC sites, to be detected by sequencing (iii-iv).

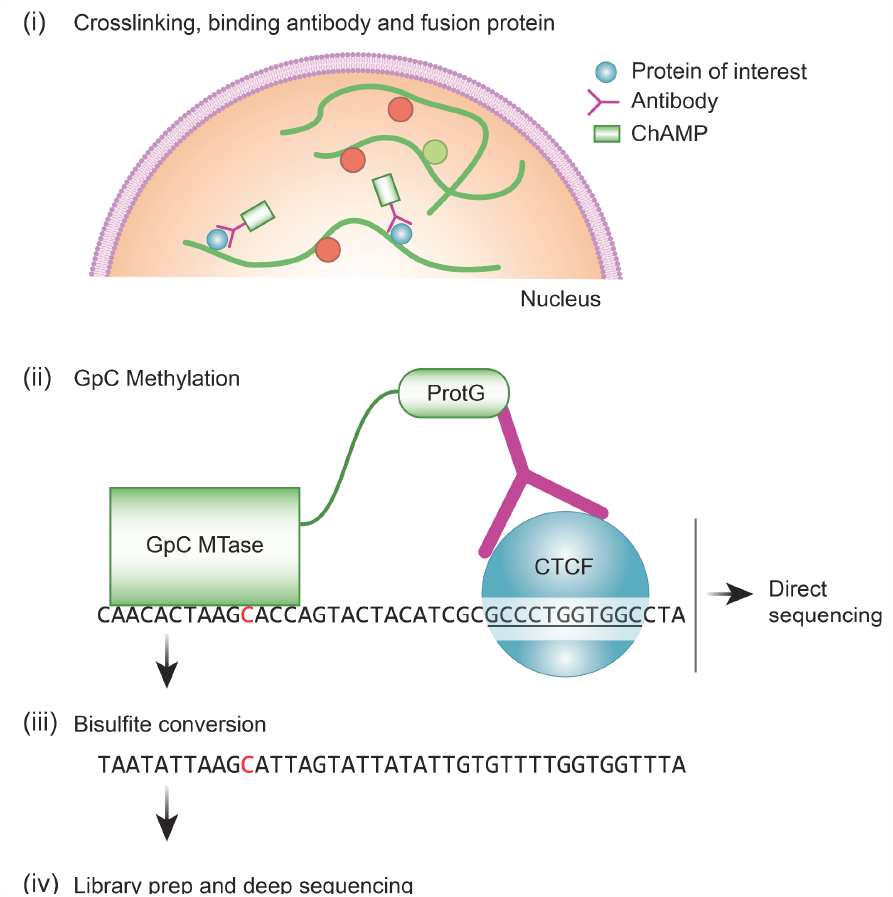

## Introduction

Characterization of protein-DNA binding sites is a fundamental aim in molecular biology. It is commonly addressed by chromatin immunoprecipitation (IP) and sequencing assay (ChIP-seq) in which the protein is immunoprecipitated, native or cross-linked to DNA, and then the bound DNA is sequenced ^1^. Challenges associated with this method include the need for an available high quality antibody to pull down the DNA-protein complex, and the loss of primary substance throughout the (IP) procedure. Hence this procedure requires large sample size (typically 10^3^-10^7^ cells) and limits the ability to analyze small samples including single cells, rare cell types and states, and limited clinical samples.

Complementary and alternative methods to ChIP-seq include DNA adenine methyltransferase identification (DamID) ^2^, Targeted Gene Methylation (TAGM) ^3^ and Chromatin Immuno/Endogenous Cleavage (ChIC/ChEC/Cut&Run) ^4,5^. DamID works by expressing an adenine methyltransferase (MTase) fused to a protein of interest, transgenically expressed in the target cells. When the fused protein binds the DNA, an adenosine methylation (which does not occur naturally in eukaryotes) will label DNA sites proximal to the binding site of the target protein. With no currently available chemistry to directly distinguish methylated adenines through clonal-amplification based sequencing methods, it typically requires specific digestion or immunoprecipitation of the methylated DNA. TAGM works in a similar way using cytosine DNA methyltransferases, overcoming the chemistry limitation. ChIC and ChEC both rely on tethering a nuclease, either by antibodies or by fusion, to a protein of interest. This results in a specific cleavage pattern, allowing for the isolation and detection of double stranded fragments from the vicinity of the binding site. Cut&Run works similarly, is performed *in situ and* without crosslinking, and is widely used to probe protein-DNA interactions. More recently, directed methylation with long-read sequencing (DiMeLo-seq) was demonstrated by targeting Hia5, a non-specific DNA adenine methyltransferase, to antibodies of interest ^6^. This enzyme was also used to observe the regulatory architectures of chromatin fibers ^7^. In a non-sequencing approach, DNA accessibility of single molecules was probed by enzymatic attachment of fluorescent cofactors, followed by optical genome mapping in nanochannel arrays ^8^.

Though adenine methylation is difficult to detect without immunoprecipitation or methylation-sensitive restriction enzymes, methylated cytosines can be detected by bisulfite conversion and sequencing. Advances in single cell methylation analysis enable reliable detection of methylation in over 50% of the genome of a single cell ^9^. But bisulfite sequencing introduces a significant reduction in mappability and causes DNA breakage resulting in short molecules. Methylated DNA can also be detected directly from single molecules using nanopore sequencing^10^.

We reasoned that antibody-mediated tethering of an MTase to a protein of interest will enable us to use antibodies capable only of binding a target of interest, while avoiding losses and inefficiencies associated with immunoprecipitation. Furthermore, an MTase with a short recognition sequence can be used to accumulate signals over a relatively short stretch of DNA, while maintaining a high mapping resolution. Specificity is provided by the fact that NpCpH methylation does not generally occur in the human genome. To achieve our goal, we developed **Ch**romatin **A**ntibody-mediated **M**ethylating **P**rotein (ChAMP) - a GpC MTase ^11^ fused to protein G. By tethering ChAMP to primary antibodies, we were able to produce distinct methylation patterns adjacent to the protein binding sites, thus facilitating their identification.

## Results

We envisioned that a protein capable of binding primary antibodies, switching-on its enzymatic activity and distinctly modifying adjacent DNA, can provide an alternative method to ChIP-seq in the characterization of protein binding sites (Graphical abstract and Fig 1A). The advantages of this approach include (i) avoidance of the sample loss inherent in the immunoprecipitation methods (ii) multiple and bi-modal adjacent DNA modifications from a single enzyme-molecule can allow for a confident detection from a single sequenced molecule (iii) the primary antibody only needs to bind to its target, and not immunoprecipitate it, enabling the use of antibodies with weaker binding capacities (iv) modifications can be directly identified from single long molecules, enabling the exploration of 3D folding and the relations between adjacent sites (v) development of new MTases with a different recognition sequence can allow the multiplexing of several DNA-binding proteins on a single molecule. This is similar to DiMeLo-seq, however, the use of GpC MTase keeps this method compatible with bisulfite sequencing.

**Figure 1.**
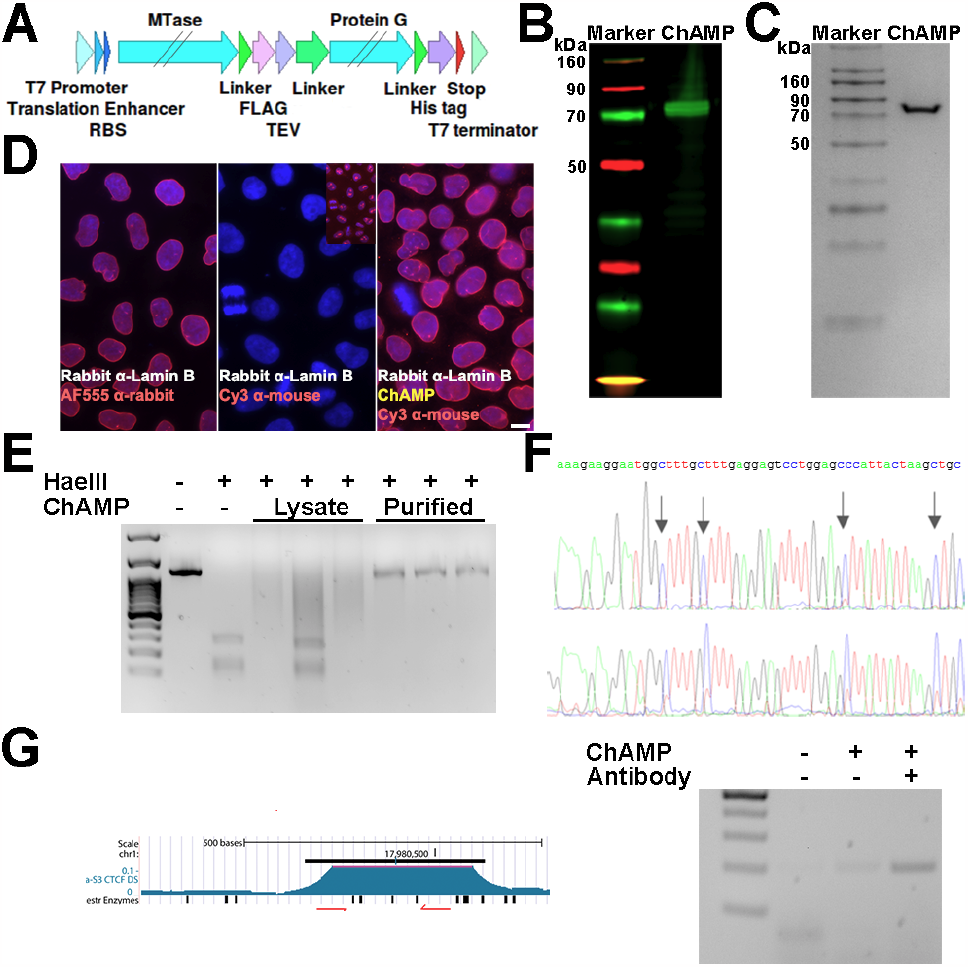
ChAMP specifically methylates GpC near the proteins of interest. (**A**) diagram of ChAMP design. (**B**) Western blot of lysed T7 Express lysY E. coli expressing ChAMP show direct binding of the secondary antibody. No primary antibody was used (see methods) (**C**) Protein staining of HIS-tag purified ChAMP. (**D**) Detecting in-situ antibody binding. Immunofluorescence of HeLa cells stained with DAPI, a primary rabbit anti lamin B antibody and (i) AF555 anti-rabbit (first panel) or (ii) Cy3 anti mouse (no binding expected - second panel) or (iii) ChAMP plus Cy3 anti mouse (bridged binding - third panel). As ChAMP has multiple IgG binding domains, it can bridge the rabbit and anti-mouse antibodies in panel 3, visualizing the nuclear envelope and proving correct localization. Scale bar - 10 µm (**E**) Detecting GpC methylation. GpC methylation inhibits HaeIII digestion. A PCR product containing multiple HaeIII cut sites was treated with either a lysate or purified ChAMP, in the presence of SAM, and digested with HaeIII. 3 purification methods were tested. Digested product was resolved on a gel. (**F**) Validation of GpC methylation in-vitro by ChAMP. Text shows genomic sequence, arrows point to GpC. Top diagram - Purified DNA was incubated with the ChAMP, bisulfite converted and Sanger sequenced. Bottom diagram - Validation of GpC methylation in-situ by ChAMP. Fixed cells were target methylated with ChAMP, DNA extracted, bisulfite converted and amplified by PCR. Significant methylation only seen upon mild fixation conditions. (**G**) Left - Restriction-PCR design overview. Genome track showing the targeted CTCF peak, HaeIII cut sites (restr enzymes) and PCR primers (red arrows). Right - ChAMP can specifically methylate DNA. Fixed cells treated with the ChAMP protocol showed a strong band following restriction-PCR, however no band at the correct size was seen when ChAMP was omitted and only a weak band when the CTCF antibody was omitted. Following the methylation reaction, DNA was digested with HaeIII and PCR reaction with limited number of cycles amplified the DNA products. Schematic representation of the method, applied to CTCF.

## Method development

We designed (Fig. 1A and supplementary data), expressed (Fig 1B) and purified (Fig 1C) a fusion protein that (a) binds to antibodies and (b) methylates GpC in the presence of S-Adenosyl methionine (SAM). We validated that the fusion protein binds to antibodies *in vitro* and *in situ* (Figs 1B and 1D) and is able to GpC methylate purified DNA *in vitro* (Fig 1E,F).

Next, we developed (Supp. Fig. 1A-C; Supp. Fig. 2; Supplementary protocol) and optimized a protocol for antibody-guided methylation of fixed samples. As a proof-of-concept, we focused on the well characterized CCCTC-binding factor (CTCF) in HeLa cells. Standard formaldehyde fixation significantly inhibited DNA methylation *in situ*. However, short fixation time with low formaldehyde concentration enabled enzyme mediated methylation (Supp. Fig. 1B). Methylation was further enhanced by a brief heating of the fixed samples prior to adding the enzyme (Supp. Fig. 1C).

**Figure 2.**
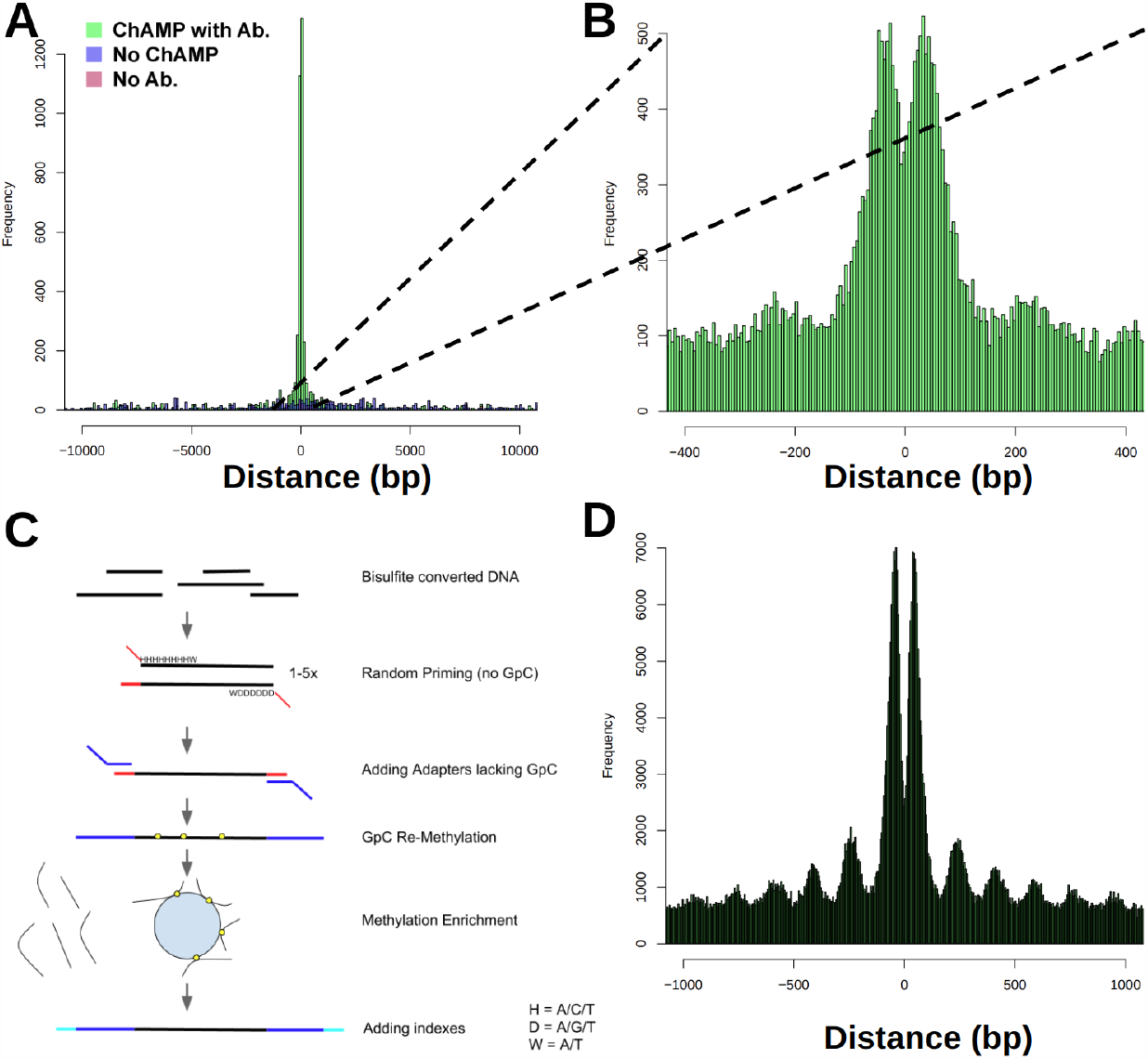
(A) Distance of dense GpCpH methylation (min. 7 events in 200bp of sliding window moving average) relative to the center of the closest CTCF ChIP-seq peak. Two experiments where ChAMP or the CTCF antibody were omitted serve as controls. (**B**) Zoom-in on ChAMP without the density filter showing a double peak. **(C)** Post-amplification enrichment **-** Schematic overview of the post-enrichment protocol. After bisulfite conversion, DNA is amplified by PCR for 1-5 rounds with random primers that do not introduce GpC. The purified product is further amplified, introducing adapters that lack GpC. The resulting product is re-methylated with the ChAMP protein. As this is amplified bisulfite converted DNA, any GpC found represents a methylated GpC (including GpCpG) in the original sample. (**D**) Distance of GpCpH methylation relative to the center of the closest CTCF ChIP-seq peak using post-amplification enrichment.

We then validated our ability to bind to primary antibodies of interest (**Fig. 1D**), to methylated CpG *in vitro* (**Fig. 1E,F**), and to *in situ* methylate near a CTCF binding site using restriction-PCR and Sanger bisulfite sequencing (**Fig. 1F,G**). While methylation was enriched near the target of interest, nonspecific binding created background methylation (Supp. Fig 3A, lower left panel). Washes with detergents minimized nonspecific binding without affecting enzymatic activity (Supp. Fig. 3A,B).

**Figure 3.**
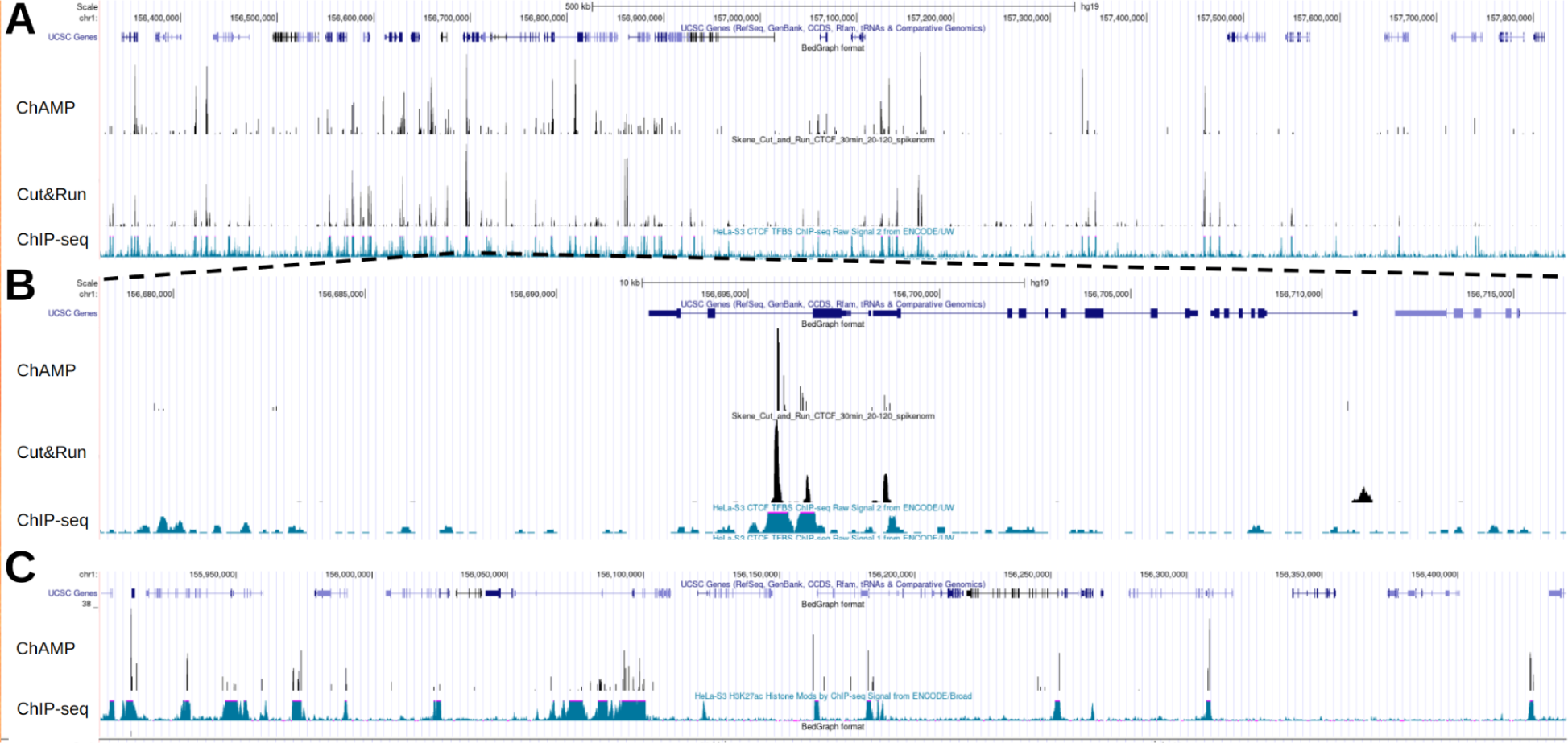
(**A**) Genome browser track showing dense methylation events for enrichment ChAMP using a CTCF antibody. These methylations partially co-localize with CTCF Cut&Run and ChIP-seq. (**B**) Zoom-in on a ChIP-seq peak. (**C**) Genome browser track showing dense methylation events for ChAMP using a H3K27ac antibody. These methylations partially co-localize with H3K27ac ChIP-seq.

### Methylation patterns

Natural GCH (H being any nucleotide but G) methylation rarely occurs in mammalian cells, and is thereby easily distinguishable from existing methylation patterns. To assess data quality, we sequenced samples to low-coverage and mapped the GCH methylation distribution relative to the closest known CTCF binding site, selecting for conditions that minimize off-target methylation. Incomplete conversion of cytosine into uracil resulted in background (typically 0.5-1%), however, as unconverted cytosines distributed randomly, requiring multiple adjacent methylation events (henceforth dense methylation) removed much of the background. To map the methylation patterns from CHAMP, we sequenced the HeLa CTCF samples to a higher coverage (122M mapped reads). We aligned all the called methylation events relative to known CTCF peak centers, then applied a sliding window threshold resulting in a strong peak overlapping the center of CTCF peaks. This pattern was not observed when either ChAMP or the primary antibody were omitted (Fig. 2A). Zooming in without the sliding window, revealed a double peak ∼30 bp on either side of the CTCF ChIP-seq peak center, presumably because CTCF and the bound antibody-ChAMP complex limit enzyme accessibility to adjacent DNA, as observed by in similar conditions ^12^. Weak secondary peaks, approximately 220bp from peak center, may be the result of nucleosome occupancy (see below).

### Methylation enrichment

Without target enrichment, ChAMP requires genome-wide coverage. While this provides the additional information of bound/unbound ratios, it requires deeper sequencing than standard ChIP-seq experiments. Enrichment for methylated DNA can reduce the sequencing requirements. However, target enrichment may involve primary substance losses, thus inhibiting our ability to apply it to small samples. We circumvented this limitation by creating a post-amplification library enrichment protocol that enriches for GpC methylation, but not CpG (Fig. 2C). In our bisulfite converted library-preparation protocol we have three consecutive PCRs, used for random priming, amplification and indexing, respectively ^9^. We modified the primers used in the first two PCR reactions to preclude the introduction of GpCs. Following the bisulfite conversion and amplification PCR, the only remaining GpCs present in the sample originated from a methylated GpC. We then re-methylated the sample (see Supplementary protocol) and immunoprecipitated the methylated DNA, effectively enriching for molecules that were originally labeled. We then performed a final amplification with indexing PCR prior to sequencing. Following the enrichment protocol, a clear structure matching nucleosome-free DNA surrounding the CTCF binding sites was seen in the cumulative data (Fig 2D). Mapping methylation sites revealed that these predominantly overlap the CTCF ChIP-seq and Cut&Run signal (Fig. 3A,B)

To validate the generality of CHAMP we tested it with an antibody directed to the histone modification H3K27ac. Dense methylated sites yielded methylation patterns that overlapped with a ChIP-seq signal (Fig. 3C). Additional methylation sites often overlapped Dnase hypersensitive regions or were adjacent to exons, suggesting the possibility of 3D folding.

### Input minimization

Reducing the minimal required cell number can allow for the exploration of rare cell types. While ChIP-seq experiments typically require a large number of cells, others have shown successful Cut&Run experiment from 1000 cells targeting CTCF^13^. By contrast, protocols for CpG methylation detection from single cells have been developed ^9^. As the output of ChAMP labeling is compatible with the input of single-cell CpG methylation protocols, we tested our ability to scale down ChAMP experiments to low (<100) cell numbers. Using the post-amplification enrichment variant, we were able to generate libraries from low cell numbers. For example, a library generated from 5 cells was able to capture 20% of all ChIP-seq peaks and showed a similar percentage overlap with Cut&Run peaks.

### Single molecule mapping of binding sites

Various single molecule sequencing technologies allow direct detection of DNA modifications ^14,15^. We used the modified version of nanopolish that is trained to call GpC methylation from Lee et al. ^10,12^. Nanopolish uses a hidden Markov model trained on enzymatically methylated DNA to distinguish between cytosine and 5-methylcytosine. While single molecule sequencing and methylation detection have higher error rates than clonal amplification and sequence by synthesis, it increases the maximal read length by 3 orders of magnitude, to over 100kb. At these scales, the extended methylation pattern resulting from a single bound site can be seen on a single molecule. Moreover, the relationship between the occupancy of neighboring sites and how distance affects it can be analysed. Despite using fixed cells, we were able to obtain multi-kilobase sequences with good quality scores (Supp. Fig. 4; N50: 8327bp). For each read, we calculate the GpC methylation probabilities for k-mers along the sequence ^12^. Methylation was greatly enriched near known CTCF binding sites, with multiple adjacent k-mers showing high probability of methylation (Fig. 4A,B).

**Figure 4.**
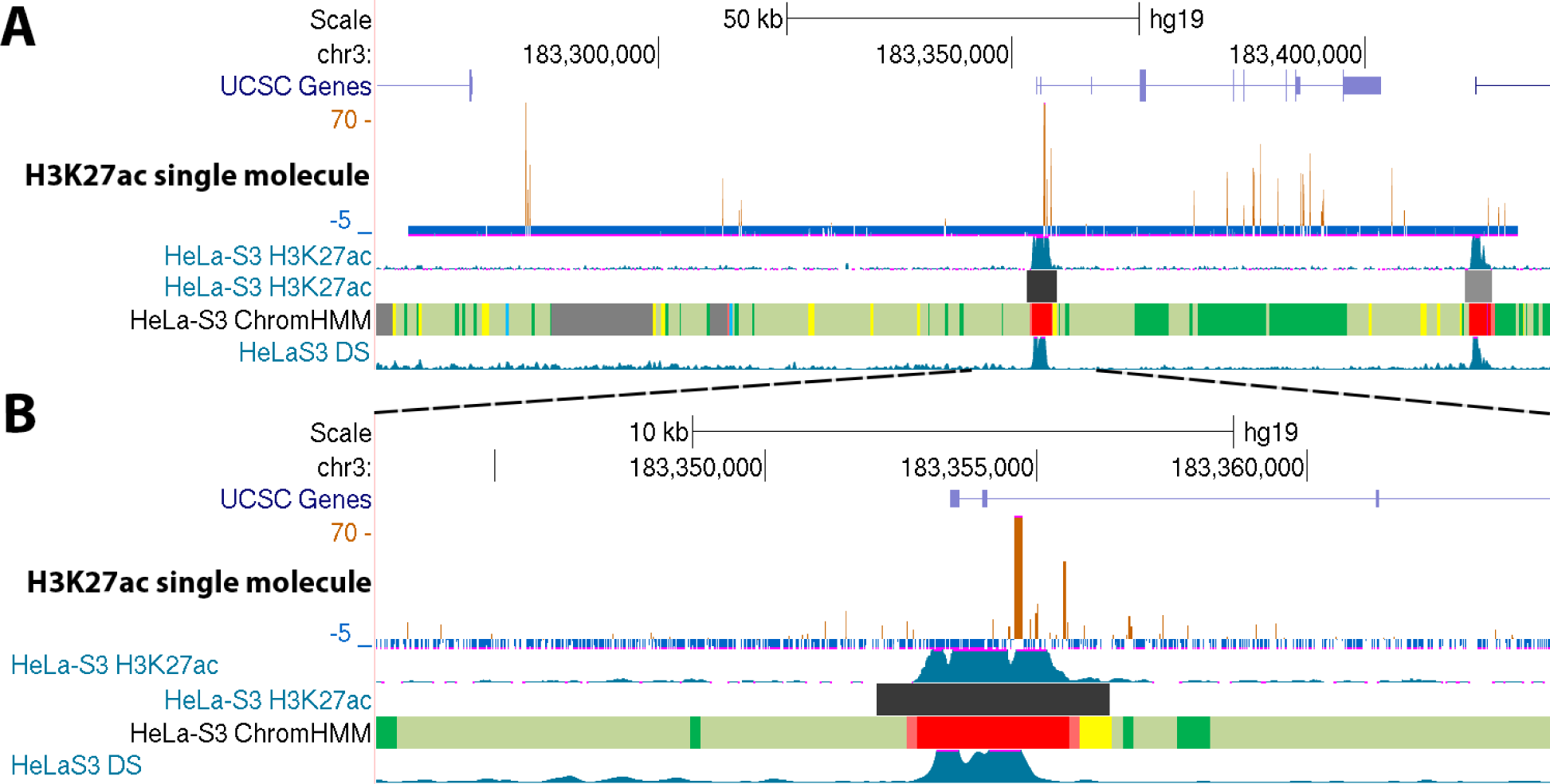
(**A**) Genome browser track showing methylation probability of k-mers along a single ∼150kb molecule from a ChAMP experiment using an H3K27ac antibody. Methylation probability was enriched, but not exclusive, to ChIP-seq peaks. (**B**) Zoom-in on a ChIP-seq peak.

## Discussion

In the current study we present a new method to map protein-DNA interactions, using low input and the possibility of mapping the binding sites on single long molecules. Importantly, the mapping is done *in-situ*, while preserving much of the DNA structure during methylation. Limitations of the system include (i) the need for a suitable antibody and motif (GpC) sites near the target of interest; (ii) the lower mappability of bisulfite-converted DNA requires more sequencing than alternative methods; (iii) ChAMP has only been tested with open chromatin near the target of interest.

But mapping protein binding on single DNA molecules can be used to answer fundamental questions about genome organization and functionality. For example, the distribution of CTCF binding sites in the genome is not random. Specifically, a ChIP-seq validated CTCF binding site has a high probability to be in the vicinity of another such site. This can be the result of sites being enriched in specific genomic regions ^17^, without any direct or indirect interactions (neutral model). Alternatively, adjacent CTCF binding sites can provide redundancy for a specific function, with only one of the sites typically being occupied (exclusive model). Finally, adjacent sites can facilitate cooperative binding and 3D organization (cooperative model). These possibilities can be distinguished by calculating how CTCF binding of one site affects the probability of CTCF binding on an adjacent site on the same molecule. This method can also allow the exploration of alleles on protein occupancy in adjacent sites.

Future studies will expand these capabilities by generating ChAMP variants targeting sequences other than GpC. A MTase lacking sequence specificity can generate more signal with improved mapping, while MTases targeting distinct sequences can be multiplexed to map the binding sites of multiple proteins on a single DNA molecule. These can even be extended to other (non 5-methylcytosine) methylations and base modifications, using single molecule technologies that can distinguish them. Moreover, direct conversion of DNA, for example by deaminases or by MTases and SAM analogs like 5’-amino-5’-deoxyadenosine, may also be used with improved genomic mapping. By directly editing DNA, all sequencing methods can be used with high accuracy and while maintaining high mappability. Development of such variants will enable the identification of multiple distinct protein-DNA interactions on a single molecule.

## Supporting information

ChAMP_v1.0

ChAMP_protocol_v1.0_Freeze.pdf

## Acknowledgments

We thank the Collins and Bar lab members for comments and suggestions. This work was supported by the Israeli Science Foundation (grants 654/20 and 632/20 to DZB), the Center for Artificial Intelligence & Data Science in Tel Aviv University (TAD to DZB) and NIH intramural support for project HG-200305 (FSC).

## Methods

### ChAMP design and synthesis

Protein DNA sequence was designed in-silico, ordered as a gblock (IDT), PCR amplified and SLICEd ^18^ into the pUC57-kan plasmid (See Supplementary file Champ_v1.0.gb). The design includes a GpC methyltransferase ^11^ fused to a protein G lacking the albumin binding domain. Linkers, a TEV cleavage site, a FLAG-tag and a His-tag were also included. Due to its methylation activity, only strains that do not restrict methylated DNA resulted in colonies after transformation. Moreover, bacteria expressing the plasmid need to be frozen or kept at the exponential growth phase, otherwise residual protein expression results in DNA methylation and bacterial death. The plasmid was maintained in 10G elite *Escherichia coli* bacteria (Lucigen).

### ChAMP purification

The full protocol is available as a supplementary file and online at link. ChAMP was expressed in T7 Express lysY *E. coli* (C3010, NEB). Bacteria were grown overnight at 37°C, 250 rpm, in 50 ml of LB supplemented with 50 µg/mL of kanamycin. Bacteria were moved into 200 ml of fresh LB supplemented with 1mM of IPTG (no kanamycin added) and grown for additional 4 hours. We note that ChAMP was readily detected without induction in both C3010 and 10G bacteria. Cells were cooled on ice for 15 minutes and centrifuged at 10,000G in a cooled centrifuge. The pellet was resuspended in GpC buffer (50 mM NaCl; 50 mM Tris-HCl; 10 mM DTT; pH 8.5) and moved to a 50ml tube. Tube was centrifuged and pellet resuspended in 10 ml lysis buffer (GpC buffer + 0.5% triton X100 + protease inhibitor + 0.1mM EDTA) and incubated at 4°C rotating for 1h. Samples were centrifuged for 10 minutes at 5,500G and the supernatant (∼10ml) was incubated with 0.5ml his-tag beads for 1h at 4°C while rotating. Beads were loaded on a column and washed with 5ml of GpC buffer followed by washes with 1.5ml of GpC buffer with increasing concentration of Imidazole. ChAMP typically eluted between 200 and 1000 mM of Imidazole, with the earlier fractions containing more protein, and the later fractions being more pure. The elutant was aliquoted to single use tubes and stored at -80°C.

### Western Blots

Samples were mixed with a protein loading buffer (Li-Cor 928-40004) and heated to 99°C for 5 minutes. Proteins were resolved on a Bis-Tris denaturing protein gel (NuPAGE, Thermo Fisher) and transferred to a nitrocellulose membrane (iBlot mini, Thermo Fisher). The membrane was blocked for 1 hour with 5% BSA at room temperature and incubated with secondary antibody (Li-Cor IRDye 925-68021 or 925-32210) for half an hour. While ChAMP contains both FLAG and His tags, no primary antibody was used, as the protein G IgG-binding domains are capable of binding most secondary antibodies directly, even after heating and membrane transfer.

### Antibody-guided methylation

The full protocol is available as a supplementary file (ChAMP_protocol_v1.0_Freeze.pdf) and online at *link*. Briefly, cells were grown on poly-lysine coated 96-well plates or T25 flasks for 24-48 hours. Samples were fixed with 0.1-1% formaldehyde for 5 minutes. Alternatively, cells were fixed by 10 minutes in cold methanol optionally followed by cold acetone for 10 minutes. Cells were permeabilized with 0.5% Triton X-100 in PBS for 7 minutes. Samples were heated to 65°C for 10 minutes and blocked with PBST with 5% BSA and 0.1M glycine for 30 minutes. Samples were incubated with a primary antibody, washed, incubated with ChAMP, washed with detergents and incubated with SAM for 1h at 37°C. Formaldehyde fixed samples were reverse-crosslinked at 65°C overnight. Proteins were digested by proteinase K and DNA used for library preparation.

### PCR validation

ChAMP methylation was validated by PCR using the primers AAGGAGAAGACAGAGTAGAGACTGC and ATGAGGAGGCTGGATGAGGT. PCR was performed using MyTaq HS Red Mix (Bioline) with 66°C annealing and 15s elongation. Real-time PCR was performed using SYBR green (Invitrogen) and annealing at 64°C. Ct values of cut were subtracted from uncut and used to calculate the fraction of uncut available for amplification.

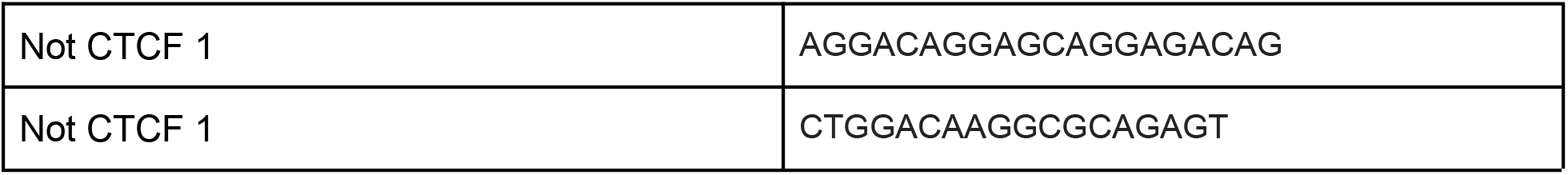
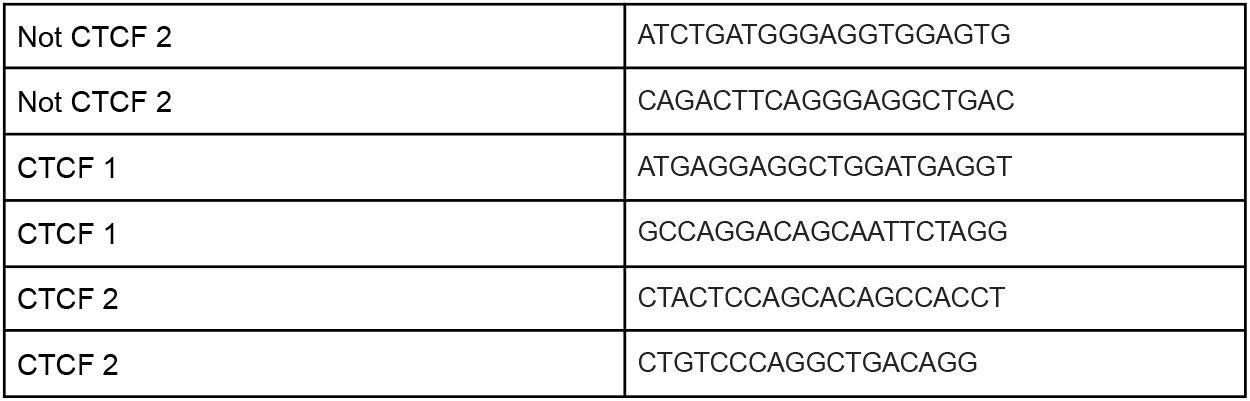

### Low input antibody-guided methylation

Resuspended HeLa cells were counted and serially diluted in medium. A single drop was placed in the middle of multiple wells in a 96 well plate, at a calculated cell number aiming for the desired range. Cells were allowed to settle in an incubator for an hour, to maximize cell adherence in the center of the well. Medium was added and the cells were incubated overnight. Two researchers independently counted the number of cells in each well. Wells with the right number of cells (2-1000 cells) were processed as in “Antibody-guided methylation”, followed by post-amplification enrichment with 5 cycles of random priming.

### Library preparation

The library preparation protocol is as a supplementary file and online at *link*.

### Nanopore libraries

Modified DNA was purified by phenol-chloroform extraction and libraries were prepared using the Rapid Sequencing (SQK-RAD004; Oxford Nanopore) kit, according to manufacturer’s instructions.

### Data availability

Sequencing data has been deposited to https://www.ncbi.nlm.nih.gov/bioproject?term=PRJNA1041053.

---

**Supplementary Figure 1.**
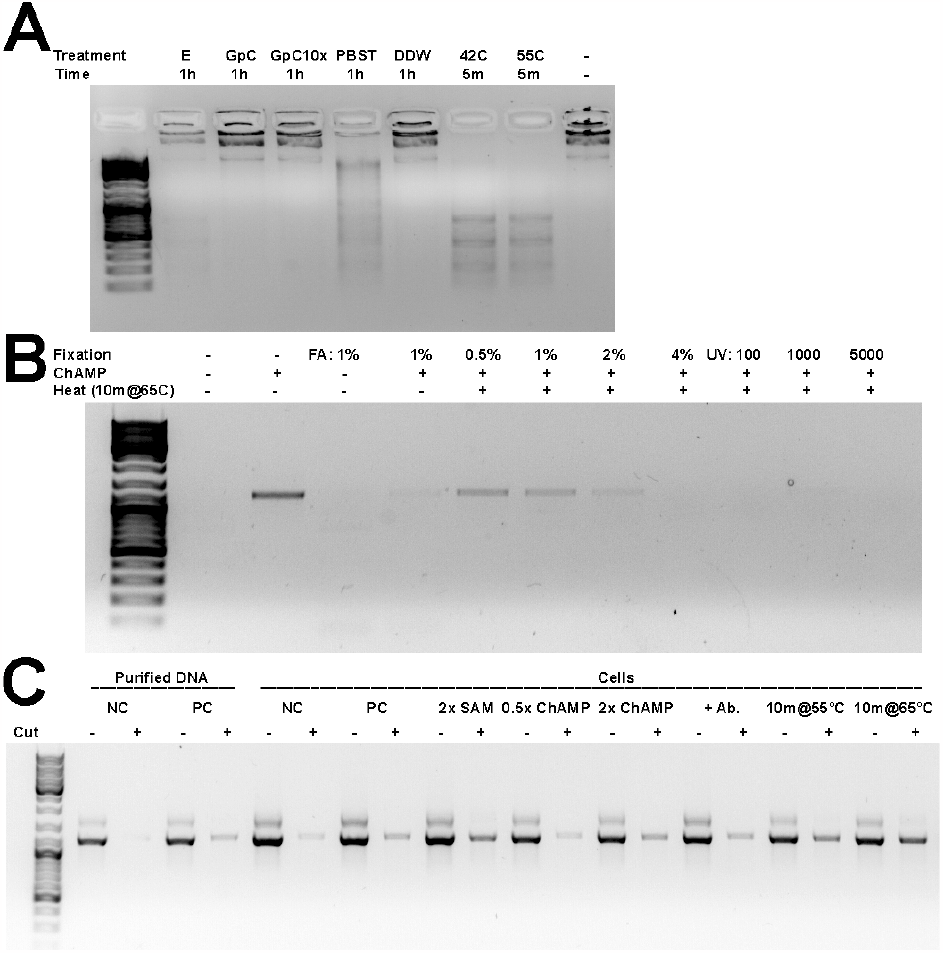
ChAMP methylation optimization. (**A**) To test the stability of ChAMP, it was pre-incubated at one of the indicated treatments (buffer/temperature/time) before being used to methylate a plasmid. Enzymatic activity was through plasmid digestion by HaeIII. E - original elution buffer, changed to GpC buffer following this experiment; no buffer indication - DDW; PBST - BPS with 0.1% tween 20; no time indication - moved directly to methylation reaction. ChAMP is sensitive to high salt and elevated temperature. (**B**) Increased formaldehyde fixation and UV exposure impairs ChAMP methylation in situ, presumably through reduces enzyme accessibility, as measured by restriction-PCR. UV crosslinking was performed using Stratagene Stratalinker 2400 UV Crosslinker at 254 nm with the indicated energy (microjoules/cm^2^ × 100) levels. (**C**) The effect of changes to the ChAMP protocol to increase DNA methylation, as measured by restriction-PCR. After methylation, DNA was digested with HaeIII prior to amplification and resolution on an agarose gel. Abbreviations: NC - negative control (no ChAMP); PC- positive control (base ChAMP methylation protocol); +Ab - CTCF antibody added (PCR product near CTCF binding site)

**Supplementary Figure 2.**
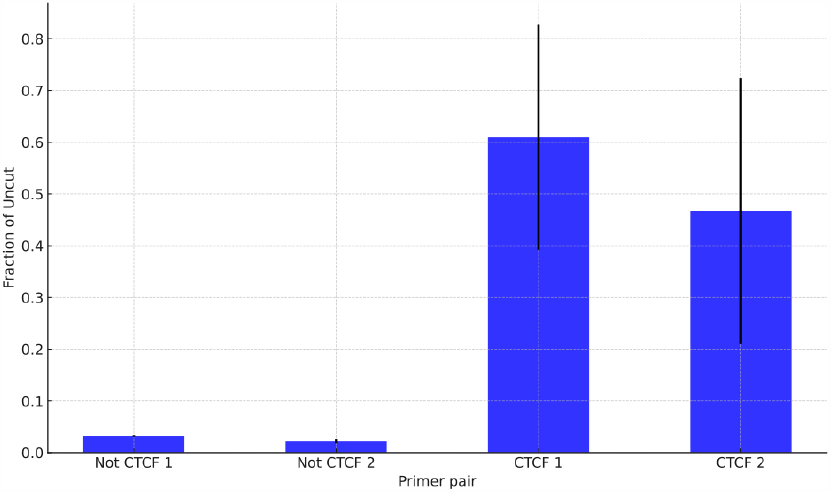
Real-time PCR validation of ChAMP methylation. After performing the full ChAMP protocol with a CTCF antibody, methylation was tested near CTCF binding sites, as well as sites not adjacent to CTCF binding sites. Primers were selected to result in a short product suitable for real-time PCR, but separated by HaeIII cut sites. Real time PCR was performed on cut and uncut DNA, and this was used to estimate the fraction of DNA available for amplification. DNA available for amplification was observed near CTCF sites, but not in control sites, suggesting ChAMP methylated specifically near these sites. Error bars represent SEM.

**Supplementary Figure 3.**
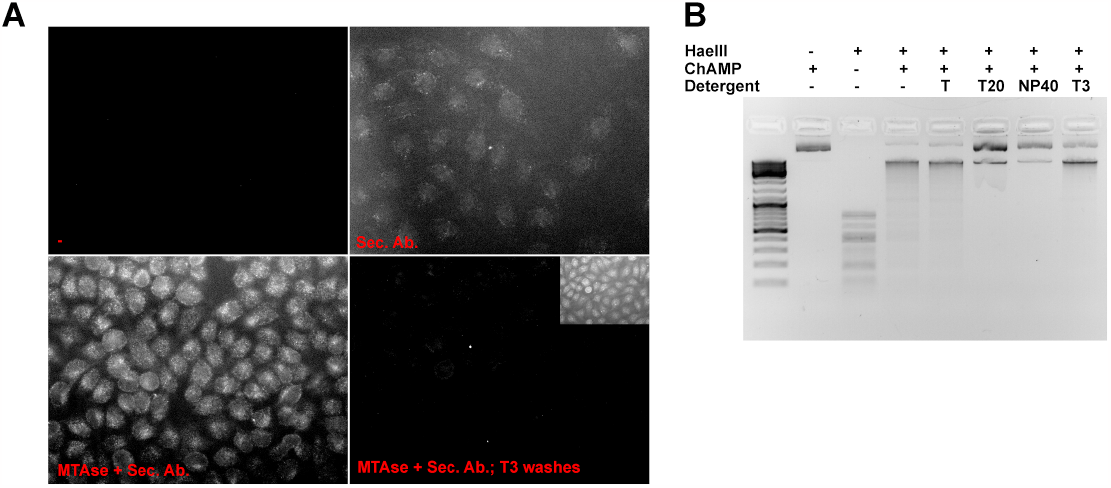
Minimizing ChAMP nonspecific binding. (**A**) Detergents reduce non-specific binding of ChAMP. Immunofluorescence images showing reduced non-specific binding of ChAMP to HeLa cells after washing with detergents (lower right vs. lower left panels). ChAMP was incubated with fixed, permeabilized and blocked HeLa cells, exposed to AF555 (Sec. Ab.) and (lower right panel) washed with detergents (T3: 1% NP40 + 1% triton + 1% tween-20). **Inlet** (lower right) - overexposure of the same field of view. (**B**) Detergents do not inhibit the enzymatic activity of ChAMP. An image of a DNA gel comparing the products of cutting a plasmid containing multiple HaeIII cut sites that was treated with ChAMP in the presence of a single detergent (1% of Triton X100 (T) or tween 20 or Nonidet P-40 (NP40)) or a combination of all three (T3; 1% each).

**Supplementary Figure 4.**
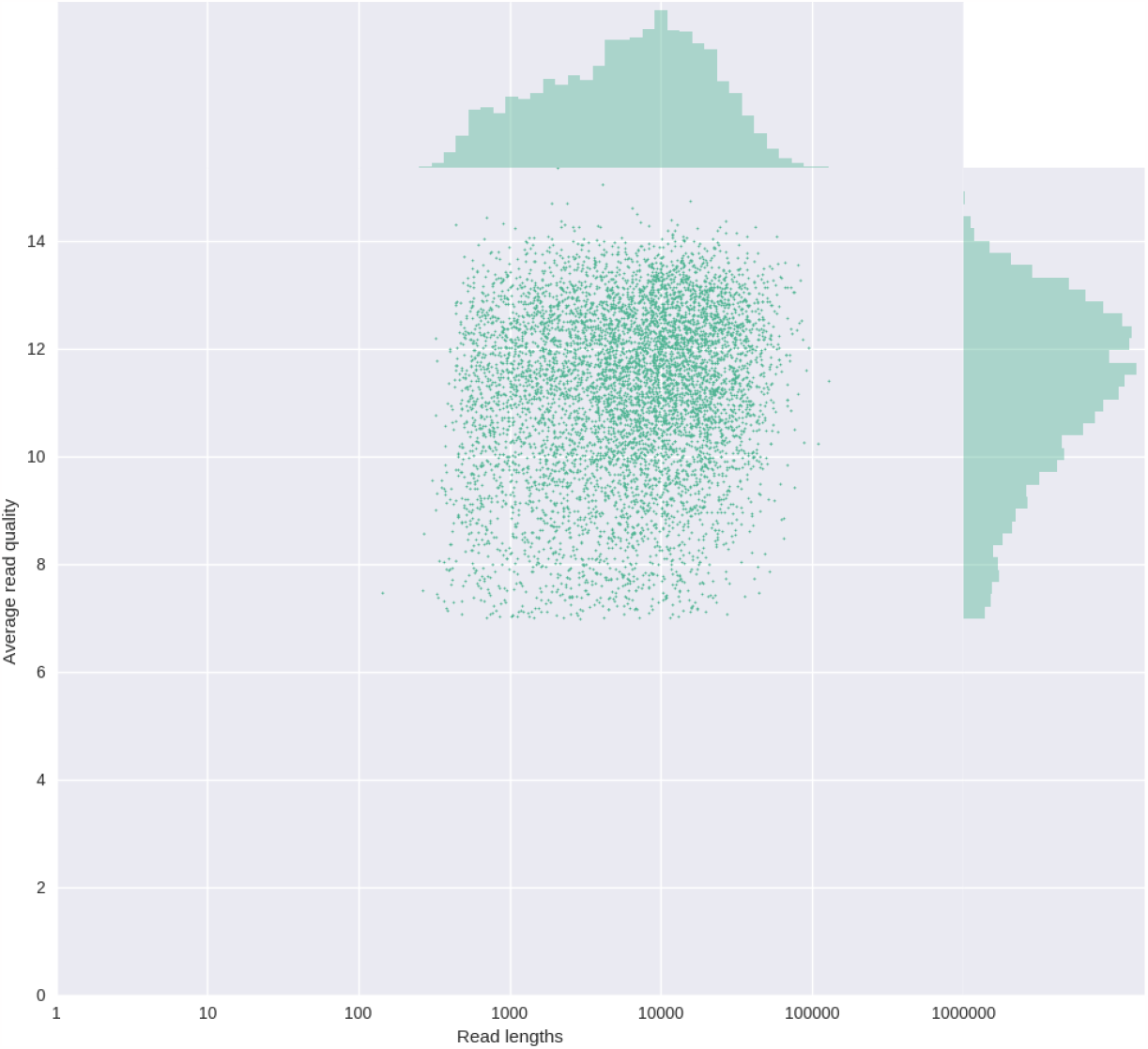
Nanopore Read length vs. quality score of a single ChAMP CTCF experiment that ran on a MinION (R9.4) cell was visualized using NanoPlot ^16^.

