## Supplementary material for "Detecting Protein-DNA Binding in Single Molecules using Antibody Guided Methylation": ChAMP_protocol_v1.0_Freeze.pdf

### ChAMP protocols

**_In-situ_ Methylation**
**Library Preparation for Nanopore Sequencing**
**Library Preparation for Illumina Deep Sequencing**
**ChAMP Purification**

#### _In-situ_ methylation

Reagents:

GpC Buffer→ 50 mM Tris-HCl, 50 mM NaCl, pH 8.5 @ 25°C. 10X stock good for years at RT.
GpC DTT → add DTT to 10mM, keep at 4C for the duration of the experiment. Good for 2 days at 4C.
PBST : PBS + 0.1% Tween 20

Recommended: Grow cells on Polylysine coated plates.
Use a T25 for direct (Nanopore) sequencing and a 96 well (10,000 cells, can be scaled down to single digit) for BSC-amplification (Illumina).

1. Wash once with PBS (NOT PBST)

2. Fix with 0.1% FA in PBST for 5mins.
   Note: Can vary according to antibody and sample - check for optimal results via immunofluorescence. Do not exceed 1% FA for 5min.

3. Immediately stop reaction with tris 1M glycine 2.5M

4. Wash once with PBST

5. Permeabilize with PBS 0.5% triton x100 for 7m
   Note: Permeabilization conditions can be adjusted depending on sample and antibody.
   Good staining by immunofluorescence is essential for successful execution of the protocol.

6. Wash twice with PBST

7. Move to 65C for 10m with PBST [50uL for 96 well, 1 mL for T25]. Make sure the plate/flask is sealed.

> Note: This improves DNA accessibility by denaturing DNA binding proteins. This will not completely remove the histones and will not affect most epitopes. Temperature can be lowered or this step can be omitted if it negatively affects immunofluorescence.

8. Block in BSA 5% (in PBST) and 0.1M glycine for 30m, Room Temp (RT) [150uL for 96 well, 1.5 mL for T25] with mild agitation. Optionally, incubate overnight 4C.

9. Incubate for 1h with primary antibody in PBST BSA 5% (1:200) at RT [150uL for 96 well, 1.5 mL for T25] with mild agitation. Optionally, incubate overnight 4C.

> Note: Primary antibody concentration can be adjusted, concentration similar to used in Immunofluorescence works well.

10. Wash 4 times with PBST over 45m. (1 quick + 3x 15min washes).

11. One quick wash with PBS + 0.5% triton.

12. One quick wash with GpC buffer.

——————————**Everything in 4C from this point on**————————

> Note: high salt concentration and elevated temperature deactivate the MTase

13. Block with cooled GpC+DTT buffer with 5% BSA for 30m [150uL for 96 well, 1.5 mL for T25] with mild agitation.

14. ChAMP mix:

> Remove ChAMP aliquot from -80C and thaw on ice (20 ng/ul stock)
> Make 1:50 dilution of ChAMP in GpC + DTT with 5%BSA

15. Aspire wells and add ChAMP premix [150uL for 96 well, 1.5 mL for T25]

16. Incubate for 1h at 4C with mild agitation.

> Note: Increase incubation time for thick samples.

17. Perform 6 washes over ~2h— all at 4C, with mild agitation:

washes 1-2: 15 mins with 1% NP40 with GpC buffer+DTT + 5% BSA + salmon DNA (1ug per well)
washes 3-4: 15 mins with 1% Tween 20 with GpC buffer+DTT + 5% BSA
last 2 washes: GpC buffer+DTT (remaining hour)

Note: Detergents needed to remove non-specific binding of ChAMP, reducing background significantly. Detergents tests as compatible with ChAMP include NP40, tween-20 and Triton X-100 and combination of all 3 at 1% each.

18. Incubate with GpC buffer+DTT with SAM 1:20 at 37C for 1h. [150uL for 96 well, 1.5 mL for T25] (SAM stock: 32mM, final conc: 1.6mM)

19. Wash twice with PBST

20. Add PBST [50uL for 96 well, 1.5 mL for T25]

21. Incubate at 65C overnight
    Note: For reverse crosslinking.

22. Add ProtK (20mgs/mL stock solution), incubate at 50C 2 hrs.
[1uL for 96 well / 5uL for T25 ]

23. Deactivate Prot K by incubating at 95C for 15 min
    Note: omit this step for direct (Nanopore) sequencing, to reduce DNA shearing.

24. Check under a microscope cells are no longer present on surface of plate [if present, add more Proteinase K + 1% SDS and continue to digest at 50C]

25. Store DNA -20C for Illumina, and 4C for Nanopore sequencing.
    Note: For Illumina, SpeedVac if necessary. Read length may decrease with storage time for Nanopore.

#### Library Preparation for Nanopore Sequencing

1. Perform a phenol-Chloroform precipitation, followed by ampure XP bead cleanup, using a 1:1 beads to input ratio.

2. Use output for library prep (either tagmentation or ligation based).

#### Library Preparation for Illumina Deep Sequencing

**1. BSC convert DNA** (based on Zymo lightning, similar kits can be used)
Add 30ng of DNA in 20ul to 130ul of lightning conversion reagent (Zymo).

Incubate in PCR

98C 8 min
54C 1.5 hours
4C <20 hours

Note: Amount of DNA can be adjusted, but should not exceed 100ng.

Add 600 µl M-Binding Buffer to a Zymo-Spin™ IC Column in a collection tube. Add the bisulfite-converted sample to the column and invert several times to mix. Spin down (max speed for 30 seconds).

Discard the flow-through and add 100 µl M-Wash Buffer to the column. Spin down.

Add 200 µl L-Desulphonation Buffer to the column and let stand for 20 minutes. After the incubation, spin down.

Discard the flow-through. Add 200 µl M-Wash Buffer to the column and spin down. Repeat this wash step.

Add 10 µl Elution Buffer directly to the column matrix and let stand for 1 minute. Spin down.

**2. Make a tube with 2ul of cleaned Bisulfite-converted DNA, 2ul of CutSmart buffer, 1ul of P5_Rand primer, 1ul of dNTP (10mM) and 13ul of DDW.**

**3. PCR 1:**

1. 98C 2 min
2. 98C 30s
3. 8C 5 min

   **Add 1ul of Klenow -exo at each 8C step, wait for temperature to get to 8C before adding.**

4. 16C 1 min

   (Ramp rate: 0.1°C/s)

5.22C 1 min
(Ramp rate: 0.1°C/s)
6. 28C 1 min
(Ramp rate: 0.1°C/s)
7. 36C 1 min
(Ramp rate: 0.1°C/s)
8. 37C 5 min
9. Goto step 2 four times
Note: 1 cycle of PCR is sufficient if greater than 100 cells
Hold at 4C

**5. Clean DNA**

Add 1ul of Exonuclease I. Incubate at 37C for 15 minutes.
Note: Remove primers.
Deactivate Exonuclease I by Incubating at 85 °C for 10 minutes.
Mix the 1.8X PCR volume of AMPure XP beads, Pipette well
Wait 5 minutes
Use magnet to collect beads and remove supernatant
Wash beads twice with 80% EtOH
Air-dry for 5 minutes.
Elute in 10ul of DDW

**6. PCR 2: Make a tube with 9ul of cleaned DNA, 2ul of CutSmart buffer, 1ul of P7_Rand primer, 1ul of dNTP (10mM) and 6ul of DDW**

Repeat PCR 1.
**Note:** Do a single cycle for greater than 100 cells, 5 cycles for less.

**7. Clean DNA**

Add 1ul of Exonuclease I. Incubate at 37C for 15 minutes
Deactivate Exonuclease I by Incubating at 85 °C for 10 minutes.
Mix the 1.8X PCR volume of AMPure XP beads, Pipette well
Wait 5 minutes
Use magnet to collect beads and remove supernatant
Wash beads twice with 80% EtOH
Air-dry for 5 minutes.
Elute in 10ul of DDW

**8. PCR with P5_handle/P7_handle (10 cycles) using Bioline Red Readymix**

Add to the 10 ul of eluted DNA

10ul X2 MyTaq HS red mix or Zymo mix
1ul P5_handle
1ul P7_handle

PCR- Red mix

95C 3m
95C 30s
60C 30s
72C 20s
Goto (95C 30s) 9 times.
72C 5m

**9. Clean DNA**

Add 1ul of Exonuclease I. Incubate at 37C for 15 minutes
Deactivate Exonuclease I by Incubating at **65 °C** for 10 minutes. *keeps DNA double-stranded
Mix the 1.8X PCR volume of AMPure XP beads, Pipette well
Wait 5 minutes
Use magnet to collect beads and remove supernatant
Wash beads twice with 80% EtOH
Air-dry for 5 minutes.
Elute in 10ul of GpC buffer
Measure using qubit (optional)

**10. Re-methylation**

Methylate DNA by adding 2ul of x10 GpC buffer (with DTT), 1ul SAM and the 3ul ChAMP and 5uL of GpC buffer, incubate at 37C for 1h. Incubate for 5m at 65C.

**11. Enrich for methylated DNA**

Enrich for methylated DNA for example using Methylated-DNA IP Kit (D5101) by Zymo:
Add 25ul of DNA Denaturing Buffer to a final volume of 50 µl.
Incubate at 98˚C for 5 minutes.
While the DNA is being denatured, mix in a 1.5 ml microcentrifuge tube:

250 µl MIP Buffer
15 µl of ZymoMag Protein A
2 µl Mouse Anti-5-Methylcytosine

Invert the tube 2-4 times to mix the antibody/Protein A mixture.
Add the denatured DNA immediately to the antibody/Protein A mixture after the incubation above is complete.
Incubate at 37˚C for 45 min on a shaking heat block.

Place tubes on a magnetic tube rack, allow time for the beads to cluster, then remove and discard the supernatant. 6.
Wash twice with 500 µl of MIP Buffer.
Wash with 500 µl of DNA Elution Buffer.
Wash with 250 uL of DNA Elution Buffer.
Add 15 µl of DNA Elution Buffer to each tube and resuspend the beads by gently flicking the tube or pipetting up and down. Transfer to a PCR tube and incubate at 80˚C for 5 minutes. Spin down and transfer the supernatant to new tube.
Measure using qubit, calculate enrichment ratio. (optional)

**12. PCR with P5/P7 (half 10 cycles/ half 20 cycles).**

Cycle number can be adjusted, depending on input DNA.
PCR-Red mix
95C 3m
    95C 30s
    60C 30s
    72C 20s
Goto (95C 30s) 9 or 19 times.
72C 5m

**12. Run on a bioanalyzer/gel**

PCR: Will notice a smear for the library- should be aiming for the library with the majority of the band between 300-500 bp.
Bioanalyzer: Will notice a more elongated curve that stretches out between 300-500bp.
Note: PCR is usually sufficient to indicate good library.

**13. Clean up and quantify:**

Mix the PCR reaction with 1.8X volume of AMPure XP beads, pipette well
Wait 5 minutes
Use magnet to collect beads and remove supernatant
Wash beads twice with 80% EtOH
Air-dry for 5 minutes.
Elute in 10ul of DDW
Measure DNA concentration by nanodrop or qubit.

**14. Proceed to deep sequencing.**

**Primers:**

| | |
|---|---|
| P5_rand | ACACTCTTTCCCTACACGACCCTCTTCCGATCTHHHHHHHW |
| P7_rand | GGTGACTGGAGTTCAGACGTGTCCTCTTCCGATCTDDDDDDDW |
| P7_handle | GGTGACTGGAGTTCAGACGTG |
| P5_Handle | TTCCCTACACGACCCTCTTCCGATCT |
| P5 | AATGATACGGCGACCACCGAGATCTACACTCTTTCCCTACACGACGCTCTT |
| P7_1_L | CAAGCAGAAGACGGCATACGAGAT**CGTGAT**GTGACTGGAGTTCAGACGTGTGCTCTTCCGATCT |
| P7_2_L | CAAGCAGAAGACGGCATACGAGAT**ACATCG**GTGACTGGAGTTCAGACGTGTGCTCTTCCGATCT |
| P7_3_L | CAAGCAGAAGACGGCATACGAGAT**GCCTAA**GTGACTGGAGTTCAGACGTGTGCTCTTCCGATCT |
| P7I1 | CAAGCAGAAGACGGCATACGAGATATCACGGTGACTGGAGTTCAGACGTGTG |
| P7I2 | CAAGCAGAAGACGGCATACGAGATCGATGTGTGACTGGAGTTCAGACGTGTG |
| P7I3 | CAAGCAGAAGACGGCATACGAGATTTAGGCGTGACTGGAGTTCAGACGTGTG |
| P7I4 | CAAGCAGAAGACGGCATACGAGATTGACCAGTGACTGGAGTTCAGACGTGTG |
| P7I5 | CAAGCAGAAGACGGCATACGAGATACAGTGGTGACTGGAGTTCAGACGTGTG |
| P7I6 | CAAGCAGAAGACGGCATACGAGATGCCAATGTGACTGGAGTTCAGACGTGTG |
| P7I7 | CAAGCAGAAGACGGCATACGAGATCAGATCGTGACTGGAGTTCAGACGTGTG |
| P7I8 | CAAGCAGAAGACGGCATACGAGATACTTGAGTGACTGGAGTTCAGACGTGTG |
| P7I9 | CAAGCAGAAGACGGCATACGAGATGATCAGGTGACTGGAGTTCAGACGTGTG |
| P7I10 | CAAGCAGAAGACGGCATACGAGATTAGCTTGTGACTGGAGTTCAGACGTGTG |
| P7I11 | CAAGCAGAAGACGGCATACGAGATGGCTACGTGACTGGAGTTCAGACGTGTG |
| P7I12 | CAAGCAGAAGACGGCATACGAGATCTTGTAGTGACTGGAGTTCAGACGTGTG |

#### ChAMP Purification

Reagents:
GpC buffer: 50 mM NaCl; 50 mM Tris-HCl; 10 mM DTT; pH 8.5
Lysis buffer: GpC buffer + 0.5% triton X100 + 0.1mM EDTA + protease inhibitor [1 tablet of roche cOmplete tablets EASYpack/50ml]

1. ChAMP was expressed in T7 Express lysY E. coli (C3010, NEB).
   Note: It is essential to use bacteria that do restrict methylated DNA

2. Grow ChAMP bacteria overnight at 37C shaker in 50mLs of LB+ kanamycin (50 µg/mL).

3. Move 10mL of overnight grown culture into 200mLs of LB+ 1mM IPTG (no kanamycin) and grow for additional 4hrs at 37C shaker.
   Note: ChAMP is readily detected without induction in both C3010 and 10G bacteria

4. Move cells into centrifuge bottles and cool on ice for 15 mins.

**——————————Everything at 4C or on ice from this point on————————**

5. Pellet cells in cooled centrifuge (4C) at 10,000 G for 15 mins. Remove media.

6. Resuspend pellet in cold GpC buffer.

7. Pellet cells again in cooled centrifuge (4C) at 10,000 G for 15 mins. Remove supernatant.

8. Resuspend pellet in 10mLs of cold lysis buffer and transfer to 50ml tubes.

9. Incubated at 4C rotating for 1h.
   Note: Sonication may also be used here.

10. Centrifuge samples for 10 minutes at 5,500G in cool centrifuge (4C).

11. Prewash 0.5ml of his-tag beads with GpC buffer, repeat once.

12. Take supernatant and incubate with his-tag beads for 1 hr at 4C rotating.

    Note: Samples of supernatant and pellet may be collected for additional Western blot QC.

13. Load a filter column with bead solution.

14. Wash filter column washed with 5ml of GpC buffer followed by washes with 2ml of GpC buffer with increasing concentration of Imidazole. Collect the passthrough in separate tubes.

    1 wash with just GpC buffer
    1 wash 20 mM Imidazole GpC buffer
    1 wash 200 mM Imidazole GpC buffer
    1 wash 1000 mM Imidazole GpC buffer

        Note: Protein eluted between 200 and 1000 mM of Imidazole, with the earlier fractions containing more protein, and the later fractions being more pure.

15. Make single use aliquots at 4C (20uL) and store right away in -80C. (Prechill tubes)

16. Quality control:

    A. Run a coomassie protein gel with the eluted fractions and a BSA standard. Successful purification will result in a strong and specific band of approximately 70kDa at the multi-nanogram/ul range.
    B. Run a DNA gel with the following reactions:
        1. 200ng of a closed plasmid #positive control
        2. Closed plasmid incubated in a reaction buffer for 1h at 37C. Digested.
        3. Closed plasmid incubated in a reaction buffer, with SAM omitted. Digested #Negative control. As HaeIII is a 4-cutter, most plasmids will have multiple cut sites.

    A successful reaction will result in only coiled, supercoiled and linear forms of the plasmid in the positive control lane, predominantly the same forms in the test lanes (increase in linear form is acceptable) and complete digestion in the negative control lane.

Reaction buffer: GpC buffer supplemented with 10mM DTT and 160 μM S-adenosylmethionine (SAM). Add cutsmart buffer and and perform function test to assess enzyme concentration, purity and function.

Digestion reaction: Add 10X cutsmart buffer directly to the methylation reaction for a final concentration of 1X. Add 1ul of HaeIII. Incubate at 37C for 1 hour.

17. After the stock has passed quality control, it can be used without re-testing for at least 6 months, as long as it is kept at -80C.
